# A breast tissue-specific epigenetic clock provides accurate chronological age predictions and reveals de-correlation of age and DNA methylation in tumor-adjacent and tumor samples

**DOI:** 10.1101/2025.02.21.639537

**Authors:** Leonardo D. Garma, Sonia Pernas, David Vicente Baz, Rosario García Campelo, Josefa Terrasa, Ramón Colomer, Desirée Jiménez, Ruth Vera García, Begoña Bermejo, Santiago González Santiago, Miguel Quintela-Fandino

**Affiliations:** Breast Cancer Clinical Research Unit, Centro Nacional de Investigaciones Oncologicas (CNIO), Madrid, Spain; Institut Catala d’Oncologia (ICO)-Institut d’Investigació Biomèdica de Bellvitge; Hospital Virgen de la Macarena, Sevilla, Spain; Medical Oncology Service, University Hospital A Coruña (XXIAC-SERGAS), A Coruña, Spain; Department of Medical Oncology, Hospital Son Espases, Palma de Mallorca; Medical Oncology Division, Hospital Universitario La Princesa, Madrid, Spain; Department of Medicine, Universidad Autónoma de Madrid (UAM), Madrid, Spain; Chair of Personalised Precision Medicine, Universidad Autonoma de Madrid (UAM – Fundación Instituto Roche), Madrid, Spain; Hospital Universitario de Fuenlabrada, Spain; Medical Oncology Department, Hospital Universitario de Navarra, Spain; Hospital Clínico Universitario de Valencia, Spain; Hospital San Pedro Alcántara de Cáceres, Spain

## Abstract

Epigenetic clocks have been widely used to estimate biological age across various tissues, but their accuracy in breast tissue remains suboptimal. Pan-tissue models such as Horvath’s and Hannum’s clocks, perform poorly in predicting chronological age in breast tissue, underscoring the need for a tissue-specific approach. In this study, we introduce a Breast Tissue-specific Epigenetic Clock (BTEC), developed using DNA methylation data from 553 healthy breast tissue samples across seven different studies. BTEC significantly outperformed pan-tissue clocks, demonstrating superior correlation with chronological age (r=0.88) and lower prediction errors (MAE=3.27 years) without requiring for dataset-specific regressions adjustments. BTEC’s chronological age predictions for tumor-adjacent samples showed distortions, with an average deviation of -1.76 years, which was even more pronounced in tumor samples, where the average difference between predicted and chronological age was -12.29 years. When analyzed by molecular subtype, the distortion was greater in the more aggressive HER2+ and TNBC tumors compared to HR+ tumors. The probes used by BTEC were associated with known oncogenes, genes involved in estrogen metabolism, cadherin binding and fibroblast growth factor binding. Despite the general rejuvenation observed in tumor tissue compared to normal breast, the correlation between BTEC’s predictions and cancer-related survival indicated that TNBC tumors with increased epigenetic ages had significant lower survival.

## Introduction

Aging is a major risk factor for numerous diseases^1,2^, and the largest risk factor for cancer^3^. As women age, their risk of developing breast cancer increases substantially, with incidence rising sharply until menopause and continuing to climb at a slower rate thereafter^4^. This age-related risk is driven by a combination of factors, including accumulated genetic mutations, prolonged exposure to hormones, and changes in breast tissue^5^.

Epigenetic changes have been found to have a strong relationship with aging^6^, and some researchers have even suggested that epigenetic modifications drive the aging process^7^. This relationship led to the development of epigenetic clocks or *epiclocks*^8–10^, which are mathematical models that predict chronological age based on the methylation state of specific regions in the DNA^11^. The difference or deviation between the predicted (or *biological* or *epigenetic*) age and the actual chronological age (i.e., the prediction error) is commonly referred to as epigenetic age acceleration (EAA) and is often interpreted as a measure of aging rate or biological aging. Epigenetic clocks have been used to link EAA to multiple features and pathologies, from cancer^1213^ to psychiatric disorders^14–16^ or socio-economic status^17,18^. However, recent research^19,20^ suggests that epigenetic clocks designed to estimate chronological age can be effectively constructed using blood DNA methylation patterns even at random CpG sites. In contrast, clocks aimed at capturing specific biological traits associated with aging would need to be based on measurement of non-stochastic, biologically regulated methylation events. Therefore, we hypothesize that accurately measuring biological age in breast tissue and uncovering the molecular changes associated with aging require the development of a novel, tissue-specific epigenetic clock. An accurate breast-specific biological aging clock could be valuable in several clinical contexts such as refining risk assessment in screening programs, predicting risk of breast cancer development or contributing to a better understanding of breast aging and hormonal influence.

Popular epigenetic clocks, such as Hannum’s or Horvath’s models, were intended as multi-organ models^9,10^. Yet, their developers observed low performances on breast tissue: Horvath reported a correlation coefficient of 0.73 between the epigenetic age calculated by its model and chronological age in 23 normal breast tissue samples of the training set of his model, and noted that the model is poorly calibrated in breast tissue. Likewise, Hannum’s clock reached a correlation of 0.72 when tested on data from 83 tumor-adjacent breast tissue samples from the TCGA, and the authors suggested that the intercept and slope of the model needed to be re-adjusted for breast. Subsequent studies testing multiple clocks reported correlations between 0.35 and 0.78, confirming the limited capability of these models to work with breast tissue data^21–23^.

Despite these limitations, these models have been applied to breast-tissue data (regressing them on age to provide a better fit) in order to address several clinical questions of relevance, such as whether DNA methylation age is or not elevated compared to other tissues in women, if there is epigenetic acceleration or not in the tumor or in the tumor-adjacent compartment compared to tumor-distant tissue, or determining whether breast tumors are epigenetically older or younger than the patient herself. To date, it has been suggested that DNA methylation age is elevated in breast tissue of healthy women^24^; that there is significant epigenetic age acceleration (EAA) in tumor-adjacent compared to normal breast tissue^22^; and that tumor tissue presents higher epigenetic age^25^ than the host.

Recently, a deep-learning pan-tissue epigenetic clock demonstrated improved accuracy in predicting chronological age when applied to breast tissue samples. However, its performance varied significantly across datasets, with r values ranging from −0.703 to 0.858 across 5 test datasets comprising 100 samples^26^. This model found that breast tumors exhibited significantly increased EAA compared to normal breast tissue, with an average acceleration of 4.542 years. Castle et al. developed a tissue-specific model with high performance (r=0.88, MAE=4.2 years) in a test set of 91 healthy donor samples^27^. While they observed no difference in EAA between normal and tumor-adjacent samples, they reported increased EAA in tumor samples. Notably, they claimed that late-staged tumors exhibited a non-significant negative EAA. Unfortunately, their model was developed using genome-wide DNA methylation data, making it incompatible with most publicly available data, which was generated using methylation microarray platforms.

Here, we evaluated the performance of various pan-tissue epigenetic clocks on a large and diverse DNA methylation dataset from breast tissue of healthy donors, consisting of 553 samples across 7 different studies. After all models produced low-accuracy chronological age predictions, we developed the Breast Tissue-specific Epigenetic Clock (BTEC), which significantly improved chronological age predictions for healthy tissue samples. When applied to tumor and tumor-adjacent samples, BTEC’s predictions were notably distorted, generally showing reduced epigenetic age acceleration (EAA) and contradicting findings from previous models. This distortion was more pronounced in tumor samples than in tumor-adjacent tissues, with the greatest effect observed in triple-negative breast cancer (TNBC) tumors, followed by HER2+ tumors, and the least in hormone receptor-positive (HR+) cases. Despite the overall decrease in EAA in tumors compared to normal breast tissue, the relationship between BTEC’s predictions and cancer survival indicated that TNBC tumors with increased epigenetic ages had significant lower survival.

## Results

### Breast tissue DNA methylation data

Previous studies examining epigenetic age in breast tissue have relied on relatively small datasets, ranging from 23 to 200^9,22,26^ subjects, limiting the robustness and generalizability of the models. This limitation is evident in the varying correlations between predicted and chronological age reported both in the original studies and in subsequent applications.

To improve upon these models, we compiled a larger dataset by integrating breast tissue DNA methylation data from multiple sources, generated using Illumina’s 450k and EPIC DNA methylation arrays. Our dataset includes samples from 13 different studies, comprising 553 normal (healthy donor) samples, 362 tumor-adjacent samples, and 1,108 tumoral samples (Figure 1A).

**Figure 1.**
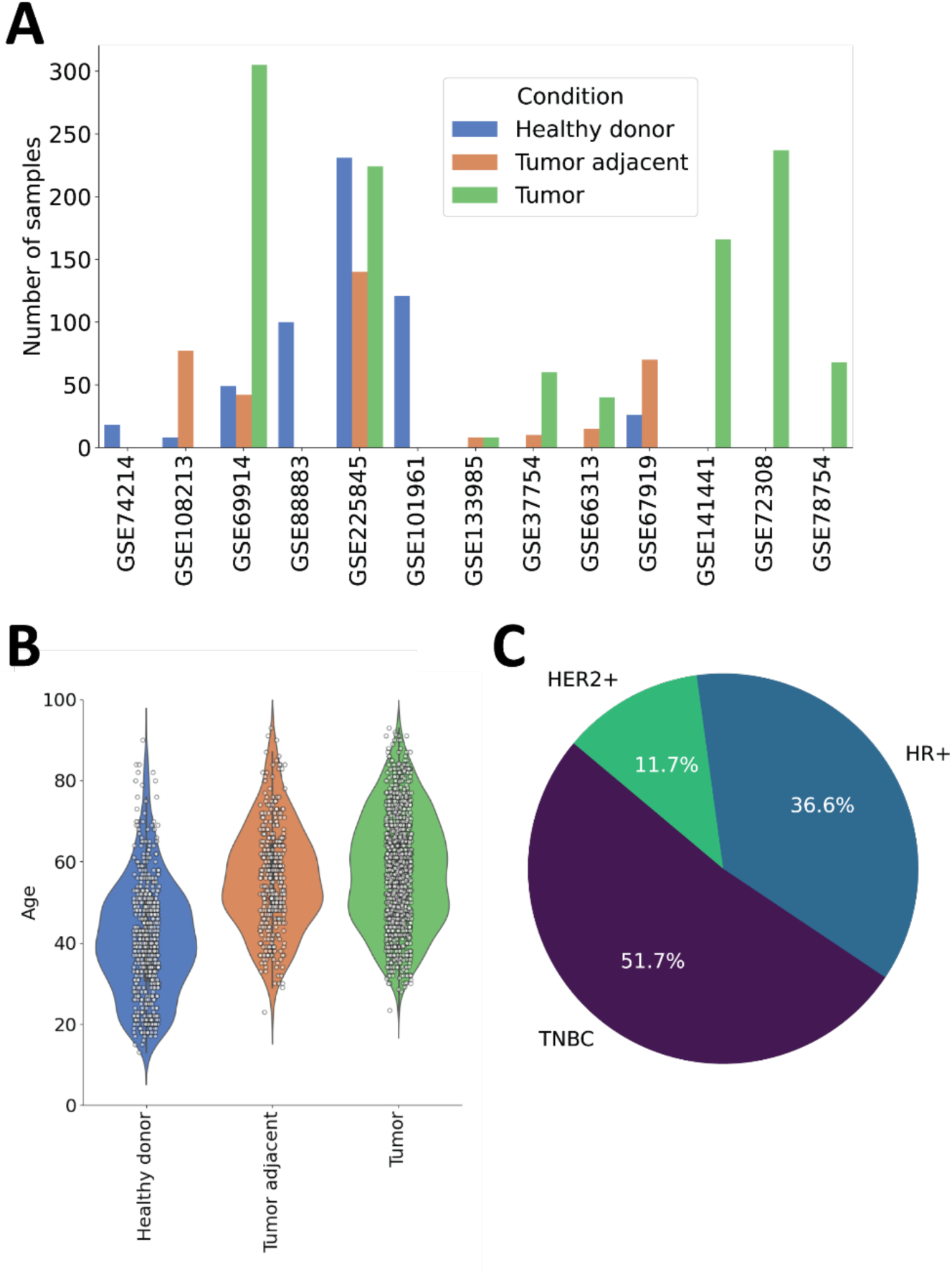
**A.** Number of samples per study and type included in the dataset. **B.** Distribution of subject ages per sample type. **C.** Distribution of tumor samples per breast cancer type.

The age of the healthy sample donors ranged from 13 to 90 years, while the tumor and tumor-adjacent samples were obtained from subjects with ages between 24 and 93 years. Out of the 1108 tumor samples, 700 were labeled with a specific molecular subtype: 256 hormone-receptor positive (HR+), 82 *HER2*-positive (HER2+) and 362 triple-negative breast cancer (TNBC) samples (Figure 1C). Detailed information about the data sources and the characteristics of the corresponding cohorts are detailed in Supplementary Table 1.

### Performance of multi-organ epiclocks

We used the combined dataset of 553 samples from healthy donors to evaluate the accuracy of chronological age predictions from four pan-tissue epigenetic clocks: Hannum^25^, Horvath^9^, PhenoAge^28^ and AltumAge^26^. Each of these models predicted the age of the donors based on breast tissue DNA methylation data (“pan-to-breast” prediction). For comparison, we two baseline models: a naïve model that always predicted the average age of the donors (41.01 years) and a random model which made random predictions within the entire age range of the dataset (between 13 and 90 years).

The results showed that the four epigenetic clocks produced predictions that were correlated with chronological age (r values ranging from 0.5 to 0.84; Figure 2A-D). However, all four models exhibited considerable root-mean squared error (RMSE) values, ranging from 9.17 to 17.58 (Table 1). AltumAge performed the best, with the lowest prediction error and highest correlation, followed by Horvath’s model. Both Hannum’s model and PhenoAge showed larger errors than the naïve model, both in terms of RMSE and median absolute error (MAE), indicating their relatively low performance. All models outperformed the random predictor.

**Figure 2.**
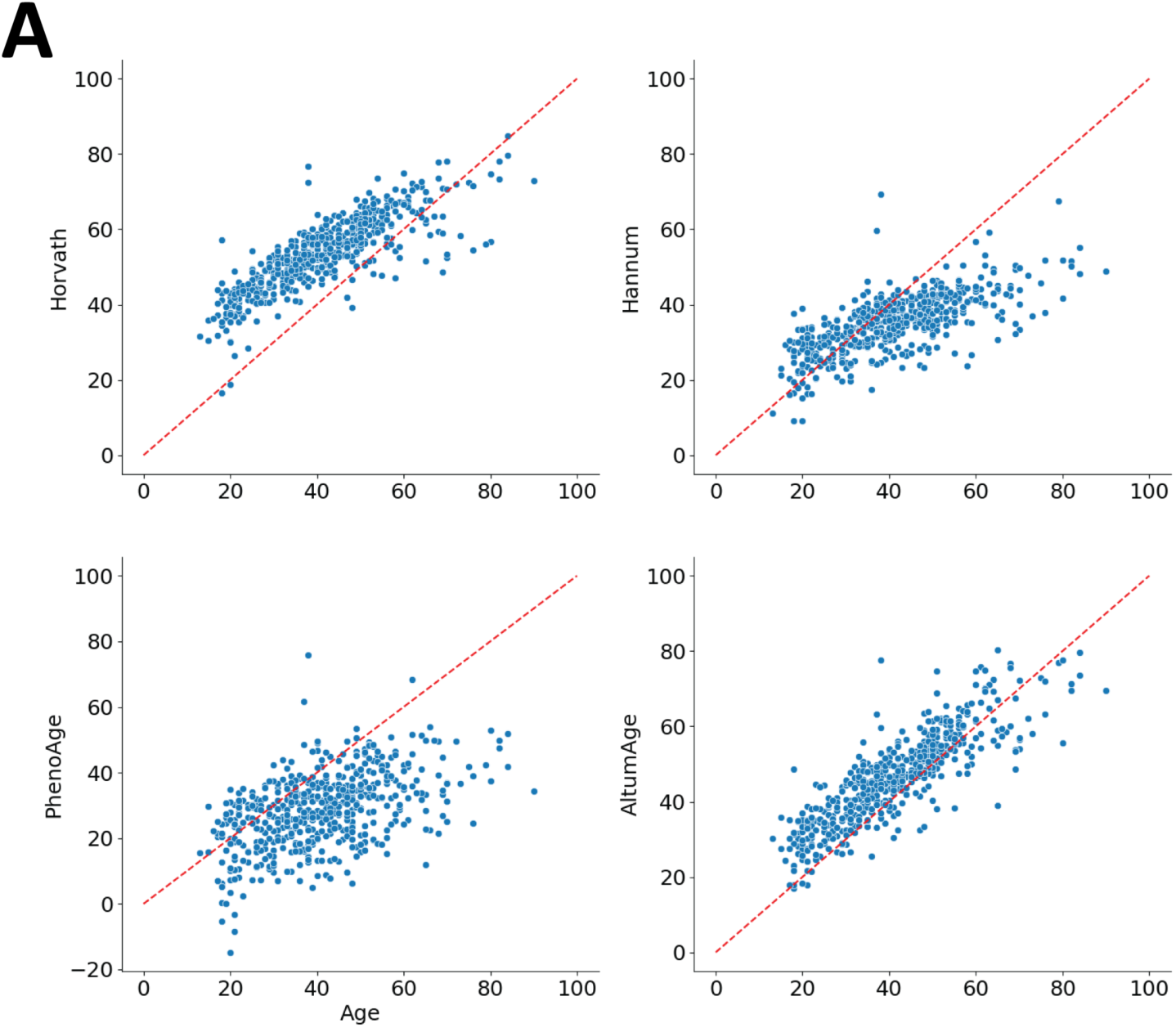
**A-D.** Chronological age predictions for the breast tissue samples from healthy donors produced by the four models tested: Horvath’s 2013 model, Hannum’s clock, PhenoAge and Altum age. Each dot indicates the actual and predicted age values of one sample and identity line is represented in red.

**Table 1.** Root-mean squared error (RMSE), median absolute error (MAE) and Pearsons’ correlation coefficient (r) for each model on the healthy donor samples.

| **Model** | **RMSE** | **MAE** | **r** |
| --- | --- | --- | --- |
| **Hannum** | 11.78 | 7.12 | 0.69 |
| **Horvath** | 14.77 | 13.63 | 0.82 |
| **PhenoAge** | 17.58 | 11.88 | 0.5 |
| **AltumAge** | 9.17 | 6.57 | 0.84 |
| **Naïve (predict the average age for all samples, 41.01 years)** | 13.93 | 9.01 | N/A |
| **Random (random values between 13 and 90 years)** | 28.10 | 20.14 | -0.06 |

### A Breast Tissue specific Epigenetic Clock outperforms pan-tissue models

To address the limitations of pan-tissue models, we developed the tissue-specific Breast Tissue-specific Epigenetic Clock (BTEC). For training this model, we utilized 406 healthy donor samples from five studies as the training set, and reserved data from two additional studies, comprising 147 samples, for validation (Supplementary Table 2).

We employed linear and quadratic terms in constructing the model, using elastic net regression. To select the appropriate CpG probes, we focused on those conserved across the 450k, EPICv1, and EPICv2 microarrays that showed significant (corrected p-value < 0.05) linear (N=201,527) or quadratic (N=179,062) correlations with chronological age in the training set. From the resulting 380,589 linear and quadratic features, we selected the top 0.5% based on the highest absolute Pearson correlation with chronological age (|r| > 0.382, N=1,962) to train the model using elastic net (Supplementary Table 3).

We set the L1 ratio to 0.5 and optimized the alpha value to 0.0013 through 5-fold cross-validation on augmented training data (see Methods). Using these optimized parameters, we applied a Leave-One-Out Cohort (LOOC) approach, where models were trained on 4 out of 5 datasets (N-1) and used to predict chronological age on the excluded cohort. This approach resulted in a strong correlation (greater than 0.89 in all cases) between the predicted and actual chronological ages. The RMSE was below 7.5 years in all but one case, where a systematic error was observed (Figure 3A, Supplementary Table 4, Supplementary Figure 1).

**Figure 3.**
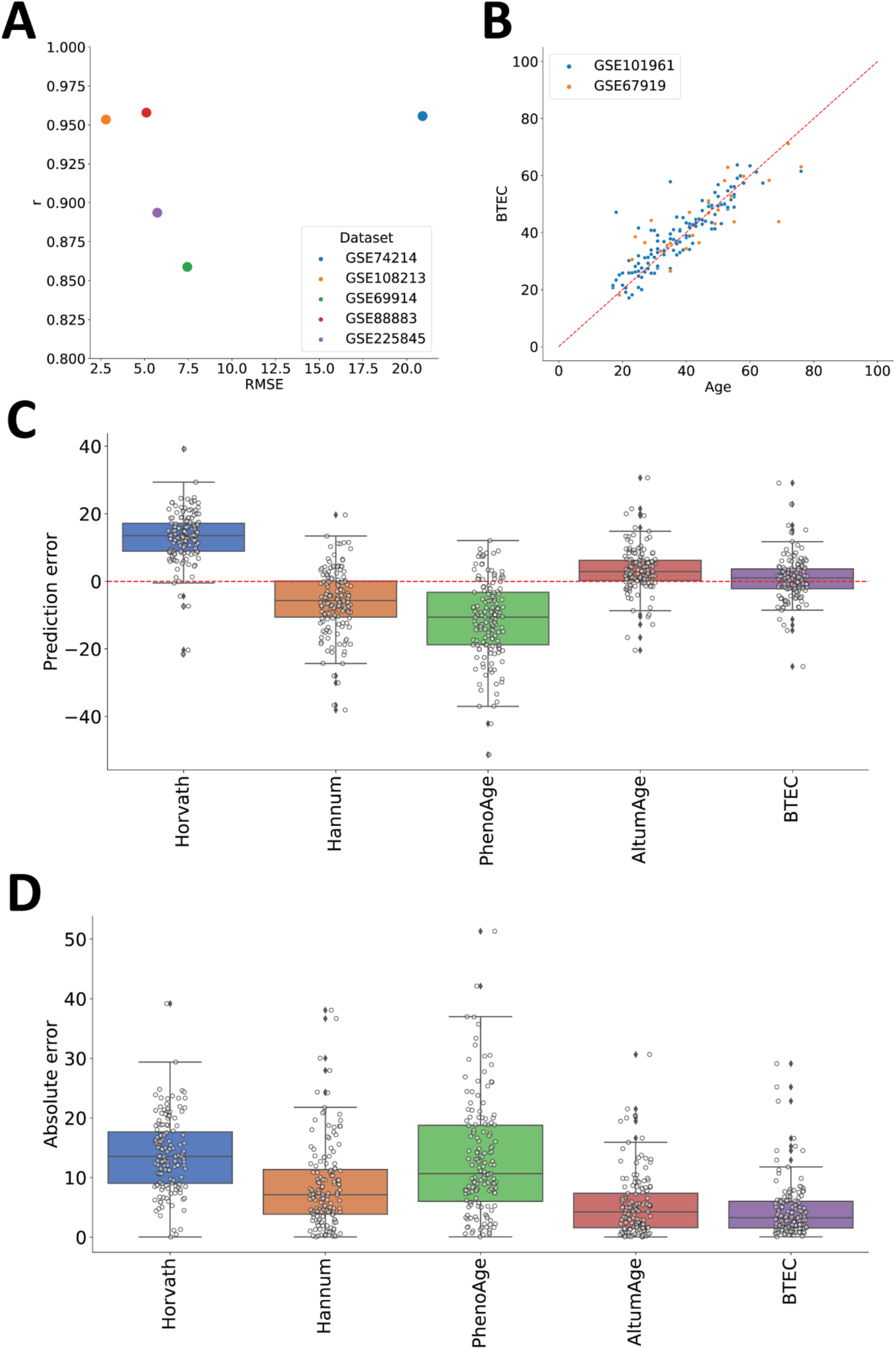
**A.** LOOC results; scatterplot of r vs MAE/MSE colored by dataset. **B.** BTEC results on validation data (scatterplot, colored by Dataset). **C.** Distribution of prediction error on validation dataset, colored by dataset. **D.** Distribution of absolute prediction error on validation dataset.

We then re-trained the model using the same parameters and the entire training set. The resulting BTEC included 637 linear and 549 quadratic terms derived from 799 CpG probes, along with an intercept value (Supplementary Table 5). When applied to the validation data, BTEC produced epigenetic age predictions that were highly correlated with chronological age (r = 0.88) and demonstrated high accuracy (RMSE = 6.31, MAE = 3.27; Figure 3B).

In the same validation data (N=147), the prediction error distribution from BTEC was the only one that did not deviate significantly from zero (one-sample t-test, p < 0.01) (Figure 3C). Consequently, the absolute prediction error was significantly lower for BTEC compared to the other four models (Figure 3D). The correlation with chronological age, along with the MAE and RMSE values indicated that BTEC outperformed the other models overall (Table 2).

**Table 2.** Results from each model on the validation dataset (N=147, healthy donor samples)

| Model | RMSE | MAE | r |
| --- | --- | --- | --- |
| Hannum | 11.09 | 7.09 | 0.71 |
| Horvath | 15.16 | 13.56 | 0.82 |
| PhenoAge | 16.03 | 10.65 | 0.54 |
| AltumAge | 7.28 | 4.26 | 0.87 |
| BTEC | 6.31 | 3.27 | 0.88 |

### Epigenetic age alterations in tumor-adjacent and tumor samples

EAA is typically calculated as the difference between the age predicted by an epigenetic clock and the chronological age. However, due to the poor performance of pan-tissue epigenetic clocks on breast tissue DNA methylation data, several studies have instead defined EAA as the residuals from a linear regression of the predicted epigenetic age on chronological age^10,21–23^. This *ad-hoc* solution introduces a bias, as the regression is computed on a per-study or per-dataset basis, making it difficult to compare results across different studies. Since BTEC produced accurate age predictions with prediction errors centered around zero for healthy tissue (i.e., it did not detect accelerated or decelerated aging in normal samples), we chose to compute EAA directly as the difference between predicted and chronological ages.

The predictions for cancer patient samples were generally lower than the chronological ages (Figure 4A). Both tumor-adjacent and tumor samples had significantly lower epigenetic ages compared to normal samples (Figure 4B). The average EAAs were −1.76 years (range: −41.74 to 30.91) for tumor-adjacent samples and −12.29 years (range: −98.86 to 56.76) for tumor samples, whereas for healthy samples, the average EAA was 0.82 years. This suggests that, contrary to other studies, BTEC’s results indicate that tumors are “rejuvenated” relative to the host. When tumor samples were categorized by molecular subtype, HER2+ samples showed significantly lower EAA (median = −18.13 years, range: −56.03 to +56.18) compared to HR+ samples (median = −4.99 years, range: −66.28 to +51.56), with TNBC cases exhibiting the largest negative EAA (median = −18.54 years, range: −71.86 to +43.8; Figures 4C and D).

**Figure 4.**
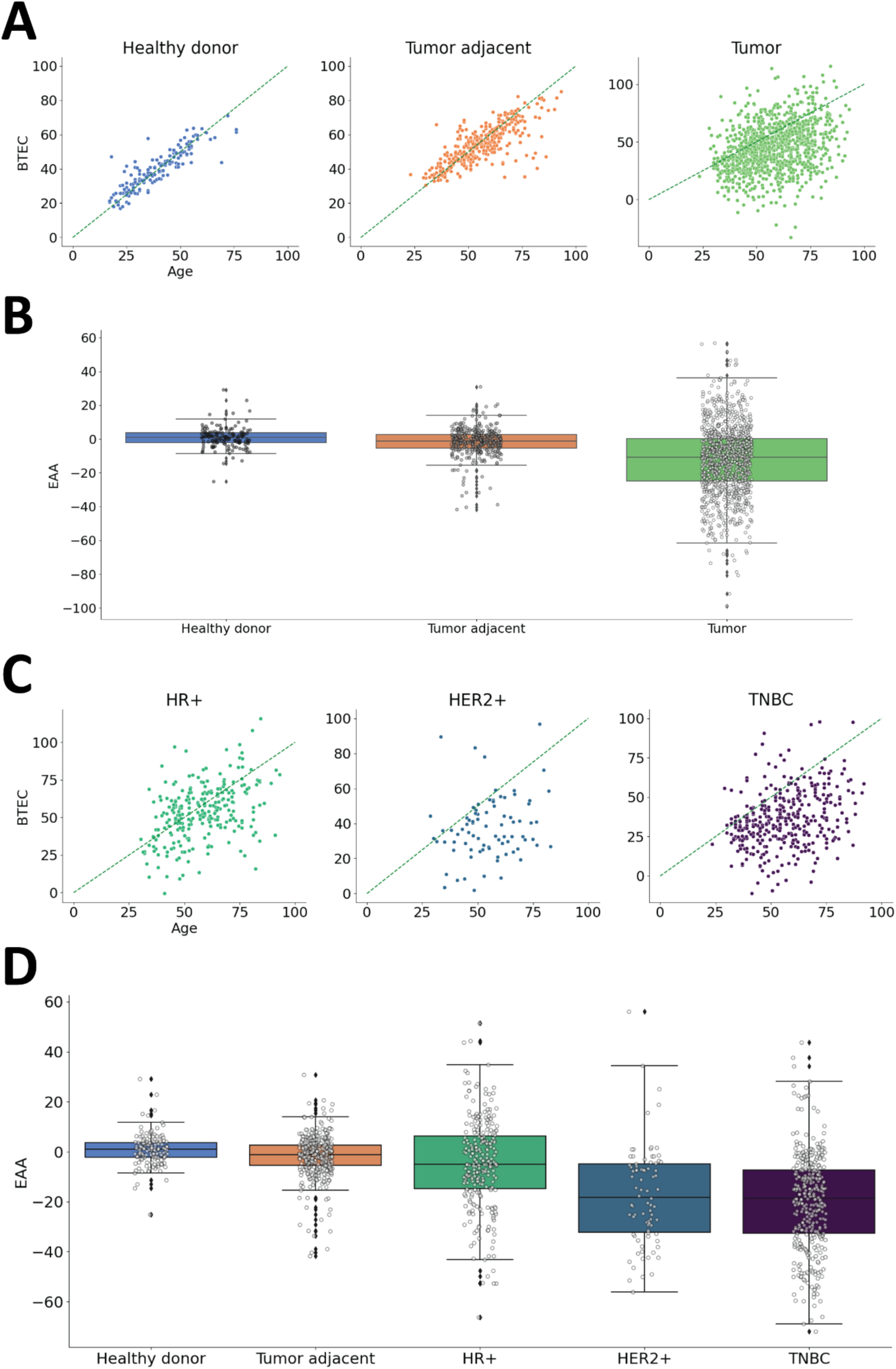
**A.** Epigenetic age predictions from BTEC on breast tissue samples from healthy donors (left), tumor adjacent samples (middle) and breast tumors (right). **B.** Distribution of epigenetic age accelerations in the three sample types. **C.** BTEC’s epigenetic age predictions on samples from hormone receptor positive (HR+), HER2 positive (HER2+) and triple negative (TNBC) tumors. **D.** Distribution of epigenetic age accelerations predicted by BTEC’s predictions broken down by tumor type.

To better understand the differences between tumors with accelerated (EAA>0) or decelerated (EAA<0) epigenetic ages as predicted by BTEC, we analyzed the transcriptomic data from Terunuma et al.’s study^29^. The differential expression analysis revealed that tumors with EAA<0 over-expressed 69 genes (p-value<0.05, two-sided Wilcoxon test, log(fold change)>1), including several collagens (*COL8A1*, *COL1A1*, *COL12A1*, *COL1A2*, *COL11A1*, *COL10A1*, *CTHRC1*) and fibronectins (*FN1*, *FNDC1*), which are involved in the epithelium-mesenchymal transition, as well as other proteins related to extracellular matrix remodeling (*CEMIP*, *MMP11*, *BGN*, *SULF1*, *FAP*) (Supplementary Table 6). These findings suggest that tumors with EAA<0 may be undergoing de-differentiation processes. In contrast, tumors with EAA>0 were enriched in 145 genes (p-value<0.05, two-sided Wilcoxon test, log(fold change)>1), including PI3k-Akt signaling kinases (*NRTK2*, *PRKAA2*, *KIT*) and genes with prognostic value (*EGFR*, *MET*, *SHC4*, *PGR*) (Supplementary Table 6). Overall, this gene expression profile indicates that tumors with EAA>0 exhibit patterns typically associated with poor prognosis.

### Tumor epigenetic age is not determined by the replication rate

Previous studies have shown that tumors exhibit accelerated biological aging^25^. To explore a possible explanation for our contrasting findings, we investigated whether the biological age of tumors predicted by BTEC correlates with the number of cell replications, as proposed by Horvath^9^. The expression of Ki-67, a well-known biomarker of cell proliferation^30^, is widely used in oncology to assess tumor aggressiveness^31^. In breast cancer specifically, Ki-67 levels are used to classify hormone receptor-positive (HR+) tumors into the Luminal A and Luminal B subtypes^32^.

Here, we used the Ki-67 annotations available in two of the datasets we collected (GSE69914^33^ and GSE141441^34^) to explore whether this biomarker was related to the chronological age of the donors, their epigenetic age predicted by BTEC, or their EAA. Ki67 data were not reported individually in the study from Gao et al.^33^; instead, patients were grouped into two categories based on their Ki-67 status: low (patients with Ki-67 below 14%) and high (patients with Ki-67 at or above 14%). This classification, based on a 13% cut-off, comes from a previous large study in which luminal-type breast cancer showed distinct clinical courses and endocrine sensitivity depending on this threshold^35^.

We observed that the distribution of chronological ages did not significantly differ between the groups (p>0.05, two-sided Wilcoxon test). However, patients with higher Ki-67 values exhibited significantly lower epigenetic ages and EAAs (p<0.05, two-sided Wilcoxon test; Figure 5A). Contrary to the assumption that accelerated biological aging is a result of repeated replication cycles, these findings suggest the opposite: tumors with lower epigenetic ages and EAAs may retain the highest replicative potential.

**Figure 5.**
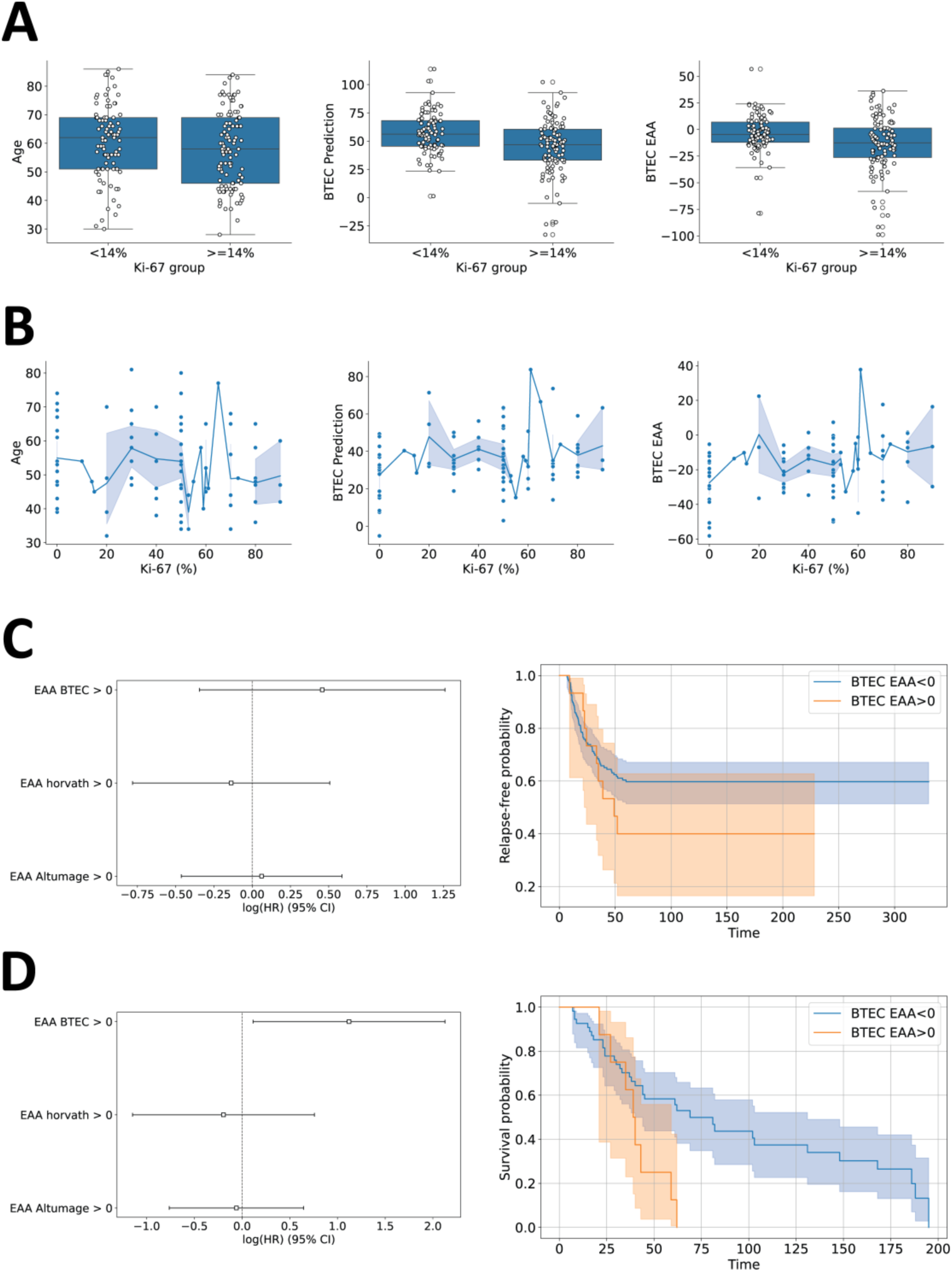
**A.** Distribution of patients’ Age (left), epigenetic age predicted by BTEC (middle) and EAA predicted by BTEC (right) on patients with Ki-67 values below or above 14% from the dataset of Gao et al.’s study^33^. **B.** Distribution of patients’ Age (left), epigenetic age predicted by BTEC (middle) and EAA predicted by BTEC (right) and their corresponding Ki-67 values on patients from Fackler et al.’s study^34^. **C.** Relapse hazard ratio for patients with EAA >0 according to BTEC, Horvath’s model or Altumage on Fackler et al.’s study (left). Kaplan-Meier relapse curves for the subjects with EAA above and below zero according to BTEC for the patients in the same study (right). **D.** Death hazard ratio for patients with EAA >0 according to BTEC, Horvath’s model or Altumage on Mathe et al.’s study^36^ (left). Kaplan-Meier survival curve for the subjects with EAA above and below zero according to BTEC for the patients in the same study (right).

In the dataset from Fackler et al.^34^, which provided detailed Ki-67 values for all subjects, we observed low correlations between Ki-67 value and age (r=0.15), epigenetic age (r=0.2), and EAA (r=0.29) (Figure 5B). When we grouped subjects using the same Ki-67 threshold as in Gao et al.’s study (14%), the only significant difference (p<0.05, two-sided Wilcoxon test) was observed in the EAA values. Specifically, the group with Ki-67≥14% exhibited higher EAA values (data not shown).

The lack of consistency and contradictory results in these two studies suggest that there is no clear relationship between the epigenetic age or the EAA determined by BTEC and the Ki-67 values. This implies that the tumors’ biological age predicted by BTEC is not directly determined by the number of replications the tumor cells have undergone.

### BTEC’s EAA magnitude is associated with long-term prognosis in TNBC

To investigate whether the tumors aging rates assigned by BTEC are linked to clinical outcomes, we analyzed data from two large datasets of TNBC patients: GSE141441^34^, which includes relapse-free times for 164 patients, and GSE78754^36^, which provides survival times for 63 TNBC patients.

Our findings showed that patients with EAA>0 based on BTEC’s predictions had an increased but statistically non-significant relapse hazard (mean HR=1.58, p>0.05 likelihood ratio test; Figure 5C). However, they exhibited a significantly lower survival probability (mean HR=3.07, p<0.05 likelihood ratio test; p<0.05 log-rank test; Figure 5D). In contrast, classifying patients by accelerated or decelerated epigenetic age using the Horvath or AltumAge models did not yield significant differences in either relapse or survival outcomes. These results suggest that, unlike pan-tissue models, BTEC’s predictions from tumor tissue may have prognostic value (Figure 5C, D).

### Biological basis of BTEC predictions

To gain insight into the differences observed in the epigenetic age predictions for healthy tissue, tumor-adjacent, and tumor samples, we examined the methylation state of the probes used by BTEC across the different sample types. Overall, the average methylation levels on the 799 probes were significantly higher in the tumor-adjacent and tumor samples compared to the healthy tissue samples (p<0.05, two-sided Wilcoxon test; Figure 6A). Upon examining individual probes, we found that 751 probes in the tumor-adjacent samples and 663 probes in the tumor samples had significantly different methylation states when compared to the samples from healthy donors (adjusted p-value<0.05, two-sided Wilcoxon test, Bonferroni corrected; Supplementary Table 7).

**Figure 6.**
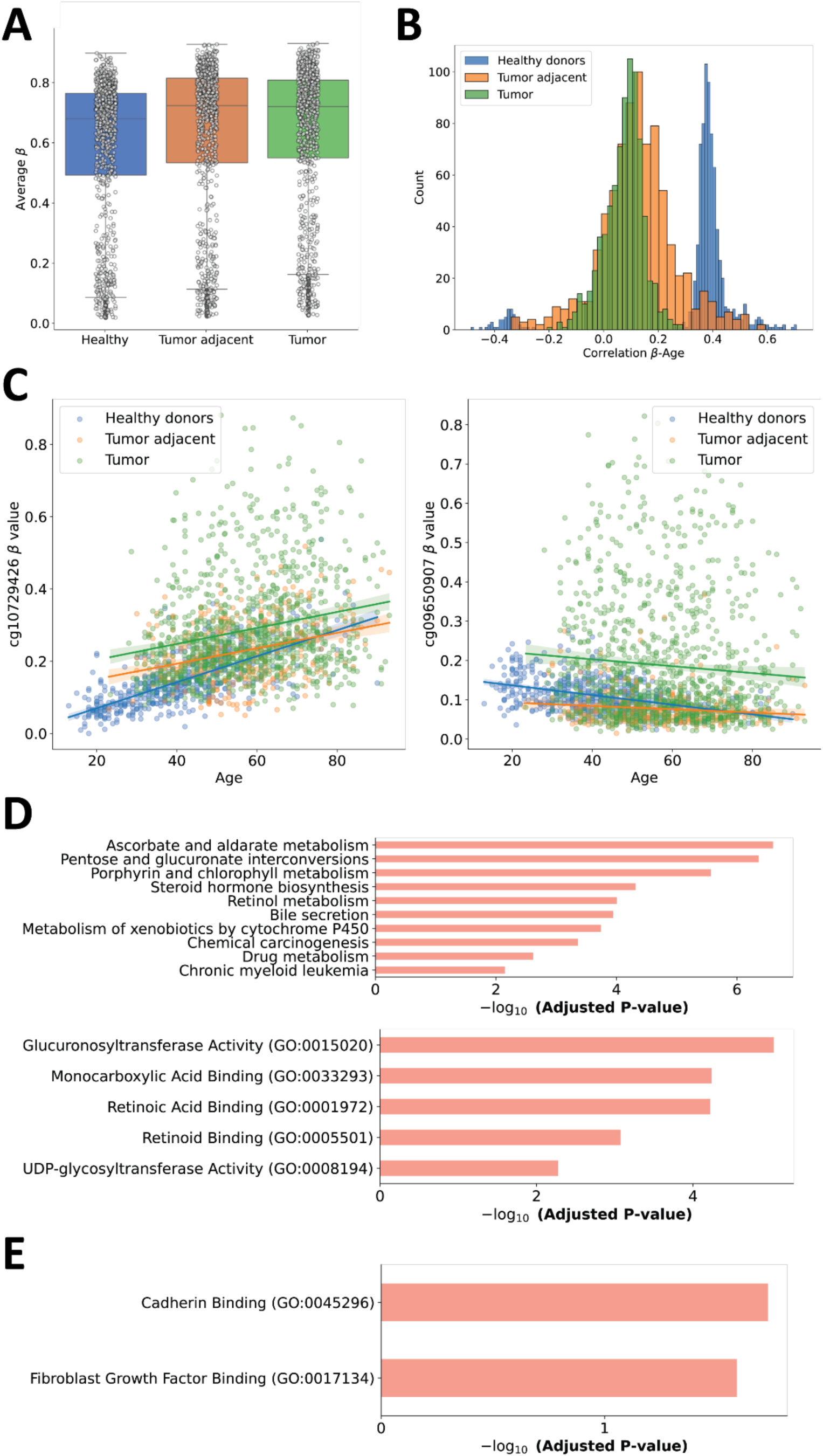
**A.** Distribution of average methylation beta values on the 799 probes used by BTEC on the three sample types. **B.** Distribution of correlations between the beta values of the 799 probes used by BTEC and chronological age, colored by sample type. **C.** Examples of probes with divergent methylation-age correlations in the different sample types. Each dot represents a sample, indicating the beta value of the probe and the subject’s age. The overlaid lines are the corresponding linear regressions. The left panel shows cg10729426, which has an r value of 0.71 in the healthy samples but only 0.32 and 0.22 in the tumor adjacent and tumor samples respectively. The right panel shows cg09650907, with r=-0.41 on the healthy donor samples and r=-0.12 and r=-0.08 on the tumor adjacent and tumor samples. **D.** KEGG (top) and GO Molecular Function (bottom) terms enriched in the genes related to the probes with positive coefficients in BTEC. **E.** GO Molecular Function terms enriched in the genes related to the probes with negative coefficients in BTEC.

Moving beyond the absolute methylation values, we observed that the relationship between the methylation states of BTEC’s probes and chronological age was very different across the three sample groups. The feature pre-selection and the regularization applied during BTEC’s training resulted in probes which had an average absolute correlation (|r|) with age of 0.4 in the healthy donor samples. However, these correlations were considerably weaker in the tumor-adjacent (mean |r|=0.15) and tumor samples (mean |r|=0.09; Figure 6B, Supplementary Table 8). In these altered tissues, the model’s probes lost the correlations with age that they exhibited in the healthy donor samples (Figure 6C). This loss of correlation likely explains the distortions observed in BTEC’s age predictions for tumor-adjacent and tumor samples.

We then explored the genes related to BTEC’s probes and their associated molecular biological functions. The 424 probes that were positively correlated with age mapped to a total of 300 genes (Supplementary Table 9). This gene set was enriched in 10 KEGG terms^37^, all of them due to the presence of UDP-glucuronosyltransferases (UGTs), except for the term “Chronic myeloid leukemia”. Enrichment in this last term was due to the presence of multiple known oncogenes in the set (TP53, MYC, AKT1, ALB1). The set of 300 genes was enriched in 5 molecular functions, all related to UGTs (Figure 5D). The deregulation of UGTs in breast tissue has a well-established connection with breast cancer, as it interferes with cell’s ability to properly manage estrogen metabolism^38,39^. Among the 300 genes in this set, only two—TP53 and nuclear receptor corepressor 2 (NCOR2)—have direct evidence linking them to aging in mammalian models, according to the GenAge database^40^.

The 376 probes negatively correlated with age mapped to 261 different genes (Supplementary Table 9). This gene set was not enriched in any KEGG terms, however they were enriched in two molecular functions: Cadherin binding and Fibroblast Growth Factor Binding (Figure 5E). These functions are known to be implicated in breast cancer^41,42^ and their deregulation could be expected in transformed tissue. Among the genes in this set, only one—ARNTL—has direct evidence of involvement in aging, according to the GenAge database^40^. These observations suggest that BTEC predominantly relies on genes with no direct link to aging and, instead, involves genes related to estrogen metabolism and cancer pathways.

Finally, we examined the RNA expression data from the dataset GSE102088, generated in the breast tissue DNA methylation study from Song et al.^43^, to identify genes involved in BTEC’s predictions that also exhibited expression changes correlated with the subjects’ age. We identified a total of 1712 genes that showed a significant and non-weak (p<0.05, |r|>0.25) correlation between expression and age (Supplementary Table 10). Among these, 24 genes were associated with probes that had positive coefficients in BTEC (Supplementary Table 10). This set of 24 genes was enriched in phosphatase binding molecular functions (GO:0019903 and GO:0051721), with notable genes such as *TP53*, *KCNQ1*, and *MFHAS1*. Another 37 genes matched those associated with probes that had negative coefficients in BTEC (Supplementary Table 10). This gene set was enriched in Cadherin Binding (GO:0045296) and Fibroblast Growth Factor Binding (GO:00171134) molecular functions, which mirrored the enrichment observed when analyzing all the genes associated with BTEC’s negative coefficients. Although the overlap between the genes associated with age in BTEC and those identified in the expression data was relatively small, as expected due to the complex relationship between DNA methylation and gene expression^44–46^, both data sources highlighted similar pathways and proteins. These included oncogenes (e.g., *TP53*, *NXN*, *TRIM59*), phosphatase binding proteins (e.g., *KCNQ1*, *MFHAS1*), cadherin binding proteins (e.g., *PAK6*, *RPL6*), and FGF binding proteins (e.g., *RPS2*, *FGFR2*). These findings suggest a connection between these molecular functions and aging in breast tissue, further supporting the relevance of BTEC’s predictions in the context of breast cancer and aging.

## Discussion

Epigenetic clocks have emerged as valuable tools to estimate biological age based on DNA methylation patterns^11,47^. However, pan-tissue epigenetic clocks have demonstrated poor performance in breast tissue: early models reported correlations below 0.75 between predicted and chronological ages and large chronological age prediction errors^9,25^; and the accuracy of state-of-the-art epiclocks exhibited a wide range of variation across datasets, with reported r values between −0.703 and 0.858 ^26^. One possible explanation for this limitation is the scarcity of breast tissue-specific DNA methylation data, which has likely hindered the development of models that accurately capture the epigenetic aging process in this tissue. Yet, despite their shortcomings, pan-tissue models continue to be widely used in studies involving both normal and pathological breast samples^21–24^.

To address the limitations of existing models, we compiled a large and diverse dataset from 13 different studies, representing the most comprehensive collection of breast tissue DNA methylation data to date. Using this dataset, we tested four pan-tissue epigenetic clocks. Our results confirmed that these models exhibited poor predictive performance and systematic errors, with some models performing worse than a naïve approach that predicts a constant age. This highlights the limitations of applying generalized epigenetic clocks to breast tissue and underscores the necessity of tissue-specific models. To overcome these issues, we developed BTEC, a breast tissue-specific epigenetic clock, which significantly outperformed all tested pan-tissue models.

BTEC provided age predictions in normal breast tissue with lower errors than any of the pan-tissue models, without the need for ad-hoc, dataset-specific regressions. The results indicate a lack of intrinsic age acceleration, in contrast with the results reported by Sehl et al.^24^. When applied to tumor-adjacent and breast tumor samples, BTEC detected a significantly decreased epigenetic age acceleration (EAA) in both conditions compared to normal breast tissue. The decreased EAA implies that unlike what was observed based on classical epiclocks^21–23^, BTEC determined that tumors are “rejuvenated” with respect to the host. BTEC identifies tumors as ‘rejuvenated’ relative to the host. While the tissue-specific model by Castle et al. indicated that epigenetic age is generally accelerated in tumors but decelerates in late-stage cases^27^, our findings show that BTEC consistently detects a strong trend toward tumor rejuvenation.

Notably, when comparing different tumor subtypes, we observed that HER2+ tumors exhibited lower EAA than HR+ tumors, and triple-negative breast cancer (TNBC) tumors displayed even lower EAA than HER2+ tumors. This suggests that tumor subtype influences epigenetic aging patterns in breast tissue.

Our analysis of the methylation states of the probes included in the BTEC model revealed that it relies on only three genes with strong evidence of involvement in aging in mammals (*ARNTL*, *NCOR2*, *TP53*). Instead, BTEC predominantly utilizes the methylation state of genes and pathways linked to breast cancer, including UGTs, known oncogenes (*TP53*, *MYC*, *AKT1*, *ALB1*), as well as FGF-binding and cadherin-binding proteins. This suggests that the aging process in breast tissue differs from patterns observed in other tissues. The distinction underscores the necessity of tissue-specific epigenetic clocks and implies that epigenetic aging in breast tissue may be more closely linked to oncogenic processes than to conventional aging pathways.

We observed that the methylation states of the sites included in BTEC were significantly altered in both tumor-adjacent and tumor tissues. Overall, methylation levels were elevated, and the expected correlations between methylation beta values and chronological age were lost. The overlap between age-correlated probes in healthy tissue and probes located within cancer-related genes likely explains BTEC’s sensitivity to disease status.

To further explore the biological relevance of these findings, we examined transcriptomic data. Although there was limited overlap between genes whose expression correlates with age and those associated with BTEC probes, our results suggest that age-related changes in DNA methylation and gene expression impact oncogenes, phosphatase-binding proteins, cadherin-binding proteins, and fibroblast growth factor (FGF)-binding proteins in breast tissue. Interestingly, BTEC’s EAA predictions were not associated with Ki-67, a widely used proliferation marker^30,31^, suggesting that epigenetic aging in breast tumors operates independently of proliferation rates.

Tumors exhibiting accelerated epigenetic age displayed transcriptomic patterns indicative of poor prognosis. In contrast, tumors with decelerated epigenetic age appeared to undergo dedifferentiation. Clinically, TNBC patients whose tumors exhibited accelerated epigenetic aging had significantly lower survival rates, suggesting that epigenetic aging profiles could have prognostic value in this breast cancer subtype.

The “rejuvenation” observed in breast tumors, as indicated by the decelerated epigenetic age and lower EAA, raises the possibility of targeted therapeutic interventions. The fact that tumors seem to undergo de-differentiation processes may suggest a potential avenue for treatments aimed at reversing such changes. Anti-aging therapies, such as senescence modulation or telomere uncapping, could hold promise in this context. Senescence is a known response to DNA damage and stress, often acting as a barrier to tumor progression. By targeting the senescent cells within tumors, it may be possible to influence their epigenetic age and potentially improve treatment outcomes. Similarly, telomere uncapping therapies could counteract the rejuvenation seen in tumors, as telomere length and maintenance are closely associated with aging and cellular senescence. Future studies should investigate how such interventions could be combined with traditional treatments to improve efficacy, particularly in the context of aggressive breast cancer subtypes like TNBC.

Given that we have demonstrated that the epigenetic age of both healthy and tumor tissue has significant value, an interesting next step would be to explore the integration of epigenetic age with other risk factors, such as breast density. Breast density has long been recognized as an important risk factor for breast cancer, but it currently remains an incomplete predictor of individual risk^48,49^. Combining epigenetic age data with breast density measurements could significantly improve risk models, providing a more accurate picture of a patient’s cancer risk profile. This could prove valuable for both early detection and personalized screening strategies. Further research is needed to determine the best way to incorporate these combined biomarkers into routine clinical practice for risk stratification and monitoring.

Taken together, our findings establish BTEC as an epigenetic clock capable of producing reliable chronological age predictions in healthy breast tissue. Its application to tumor-adjacent and tumor samples provides new insights into the epigenetic alterations associated with breast cancer. While breast cancer generally induces systemic increases in EAA, our results suggest that breast tumors and their surrounding tissue experience epigenetic changes in the opposite direction. Additionally, our findings emphasize the role of breast tissue-specific genes in the aging process and highlight BTEC’s potential utility as a prognostic tool. Future research should further explore the molecular mechanisms underlying these epigenetic alterations and assess the broader clinical implications of BTEC’s predictions.

## Methods

### Study cohorts

We used publicly available methylation data from 13 previous studies, which we retrieved from the GEO database^50,51^. We include the GEO accession number, the number of samples per condition and tumor subtype (when available), as well as the minimum, maximum, and median ages of the subjects in Supplementary Table 1.

### Data collection and preprocessing

Methylation data from previous studies were obtained from the GEO database. In all cases, the data were preprocessed: we used the beta values provided by the original authors when available; otherwise, we calculated the beta value from the methylated and unmethylated signals using the following formula:

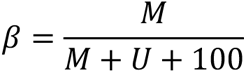

where M and U are the intensities of the methylated and unmethylated signals respectively.

We modified the original sample labels as follows: we labeled as “Tumor adjacent” all samples with the annotations “adjacent normal” and “ipsilateral normal”; samples with “reduction mammoplasty” and “prophylactic” were relabeled as “Healthy donor”; and samples annotated as “DCIS” were labeled “Tumor”. Samples labeled “controlateral” were discarded.

On the tumor samples, 4.43% of the beta values from the 366863 cross-platform (450k, EPIC, EPICv2) probes were missing in at least one samples. We imputed the missing values with the median value across all samples for each probe. On the non-tumor samples, we imputed 4.4% of the beta values using the same strategy.

### Epigenetic age prediction using existing models

We used four pan-tissue epigenetic clocks to predict chronological age using DNA methylation data from breast tissue of healthy donors using the PyAging Python library^52^: Horvath^9^, Hannum^25^, PhenoAge^28^, and AltumAge^26^.

### BTEC training

We used elastic net to train a chronological age predictor based on the beta values of cross-platform (450k, EPIC, EPICv2) probes which exhibited high correlation with age in the set of heatlthy samples. Specifically, we considered the probes with the top 0.5% r values among all those which had a significant (corrected p-value < 0.05) linear or quadratic correlation with chronological age (a total of 1,962 features; Supplementary Table 3).

We used 406 samples from healthy subjects across five datasets for training the model. To reduce the effect of the imbalance by dataset, we augmented the training data by resampling each dataset to a total of 200 samples. On the resulting 1000 samples, we optimized the alpha and L1 ratio values to 0.0013 and 0.5 respectively using a 5-fold cross-validation.

The performance of the model on the training set was then assessed using a LOOC approach, and the model then was retrained on the whole augmented training data. The resulting model, consisting of 1187 parameters (an intercept value, 637 linear and 549 quadratic coefficients), was used for the age predictions on the validation dataset as well as on the tumor adjacent and tumor samples.

### Transcriptomic data analysis

In total we used transcriptomic data from two datasets to 1) compare the transcriptomic profiles of tumors with EAA>0 and EAA<0 and 2) analyze the correlations between gene expression and age.

To compare tumors with in different EAA groups, we used the dataset GSE37754 from Terunuma’s et al. study^29^, which contained microarray expression data (Affymetrix Human Gene 1.0 ST Array) from 108 tumor samples, out of which 55 matched tumor samples from the same study with available methylation data (GSE37751). The data was mean-centered and scaled to unit variance and then the two groups were compared using a two-sided Wilcoxon test.

To study the correlation between gene expression and aging in breast tissue, we used the transcriptomic data from Song’s et al. study^43^ (GSE102088), which included 104 samples from healthy donors. We used the normalized gene expression matrix provided by the authors to determine the Pearson’s correlation coefficient of each gene and chronological age.

### Gene set enrichment analysis

In all cases, we used the GSEAPY Python library^53^ to perform gene set enrichment analysis through the Enrichr^54^ API. We included the KEGG 2021, GO Molecular Functions 2023 and GO Biological Process 2023 gene sets to perform the analysis on human genes, with no specific background and an adjusted p-value cutoff of 0.05.

### Survival analysis

The hazard ratio estimations were obtained using a Cox proportional hazard model through the lifelines Python library^55^. Kaplan-Meier survival curves were obtained using the same library.

