## Supplementary Figure 1 for "A breast tissue-specific epigenetic clock provides accurate chronological age predictions and reveals de-correlation of age and DNA methylation in tumor-adjacent and tumor samples"

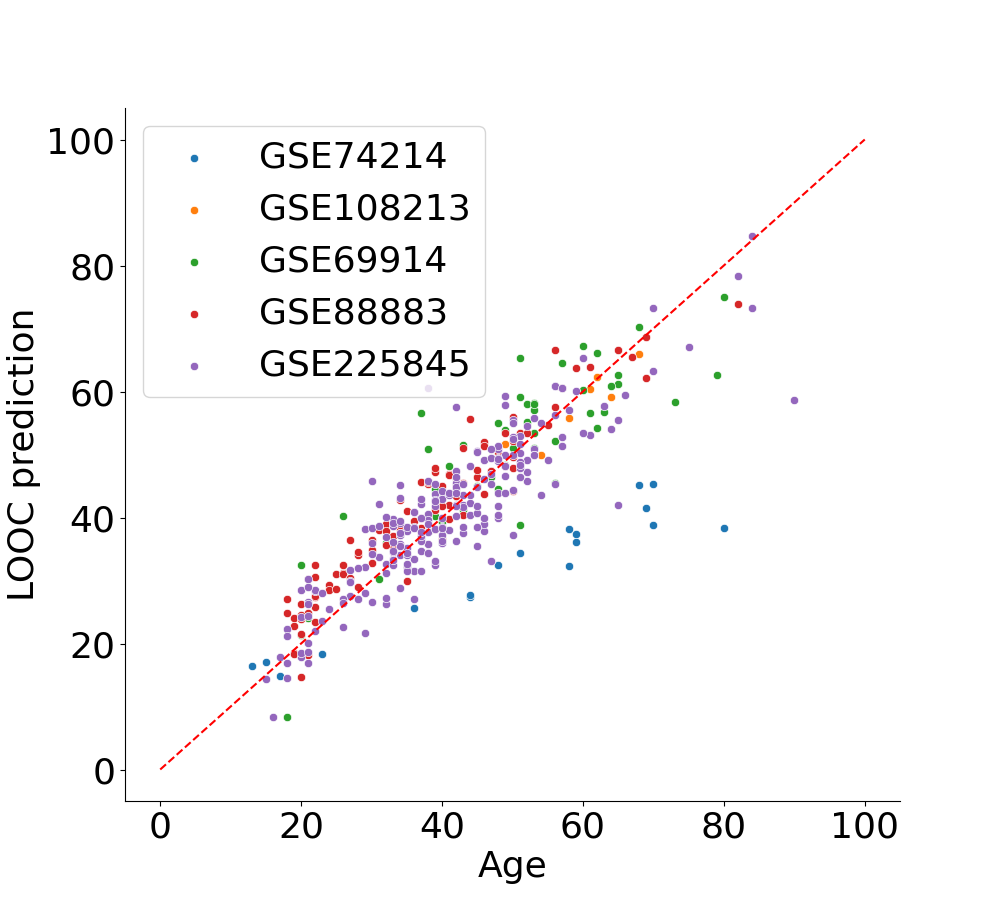

**Supplementary Figure 1.** Chronological age of donors and age predicted by BTEC during the LOOC training and assessment of the model. All datasets tested presented good correlation between predicted and actual age, however GSE74214 exhibited a systematic error, maintaining a linear correlation but deviating from the identity line.
